# A Novel NO Donor 3A72 Activates Mitochondrial-Associated Endoplasmic Reticulum Membrane Structures to Induce Ferroptosis in Triple-Negative Breast Cancer

**DOI:** 10.64898/2026.09.28.754991

**Authors:** Yunke Long, Yuxin Jiang, La Li, Xiufan Wu, Jiachen Weng, Ying Chen, Hongrui Liu

## Abstract

Triple-negative breast cancer (TNBC) remains a major clinical challenge due to a lack of effective interventions and severe drug resistance. Disrupting tumor metabolism using nitric oxide (NO) donors to drive ferroptosis represents a promising therapeutic strategy. Here, we developed 3A72, a novel furan-coumarin-based NO donor with potent anti-TNBC efficacy. *In vitro* and *in vivo* studies demonstrated that 3A72 significantly suppresses TNBC cell proliferation and restricts xenograft tumor growth. Notably, its efficacy matches the first-line drug paclitaxel but with enhanced safety and reduced systemic toxicity.

Mechanistically, 3A72 induces ferroptosis in triple negative breast cancer (TNBC) cells via a synergistic dual-pathway mechanism. Initially, 3A72 releases intracellular NO to trigger cytoplasmic calcium overload, followed by high-affinity binding to both inositol 1,4,5-trisphosphate receptor (IP R) and acyl-CoA synthetase long-chain family member 4 (ACSL4). Binding to IP_3_R provokes mitochondrial calcium overload and a massive burst of reactive oxygen species (ROS), while concurrent ACSL4 activation drives essential membrane phospholipid remodeling. Crucially, our transcriptomic and structural predictions reveal that while ACSL4 exhibits a broader distribution across organelle membranes, its functional association with IP_3_R spatially converges at the endoplasmic reticulum membranes associated with mitochondria (MAMs). This subcellular niche closely couples IP R-mediated ROS generation with ACSL4-driven lipid remodeling, accelerating lethal malondialdehyde (MDA) accumulation and triggering irreversible ferroptosis. Overall, these findings highlight the MAMs calcium–lipid metabolic hub as a promising structural scaffold for antitumor drug development.

## 1 Introduction

Triple-negative breast cancer (TNBC) is a refractory subtype of breast cancer characterized by the lack of estrogen receptor (ER), progesterone receptor (PR), and human epidermal growth factor receptor 2 (HER-2) expression[1]. Due to the absence of effective therapeutic targets, TNBC does not respond to both conventional endocrine therapy and anti-HER2 treatment regimens for breast cancer[2]; this resistance, coupled with its highly invasive clinical course, poses a formidable challenge in breast cancer management. Although novel therapeutic approaches, such as immune checkpoint inhibitors and antibody-drug conjugates (ADCs)[3], have gradually been emerged to the clinical treatment of TNBC in recent years[4,5], their efficacy is often constrained by advanced disease stage at diagnosis and a paucity of actionable molecular targets[6]. Furthermore, TNBC frequently acquires profound resistance to apoptosis during treatment[7], which consequently induces TNBC cell death via non-apoptotic pathways has become the key to breaking this bottleneck.

Ferroptosis, a distinct form of regulated cell death driven by lipid peroxidation[8], has emerged as a promising mechanism to bypass apoptotic resistance[9]. Therefore, therapeutically exploiting ferroptosis signaling pathways presents a novel paradigm for targeted TNBC treatment.

Ferroptosis fundamentally arises from the lethal accumulation of intracellular lipid peroxides. This metabolic disruption intersects heavily with mitochondria-associated endoplasmic reticulum membranes (MAMs). These 10–30 nm microdomains mediate physical contact between the endoplasmic reticulum (ER) and mitochondria[10], serving as essential hubs for inter-organelle substance exchange and energy metabolism[11]. MAMs govern calcium (Ca^2^□) homeostasis between these organelles. They also provide a vital platform for phospholipid biosynthesis, transport, and metabolic remodeling[11]. Despite their physiological importance, the structural regulation of MAMs under oxidative stress in TNBC cells remains unknown. The exact structural and spatial interactions among the core proteins controlling lipid metabolism and Ca^2^ signaling at these sites are also unexplored[12]. Uncovering the intrinsic link between MAMs integrity and TNBC proliferation is therefore critical. Specifically, deciphering how MAMs homeostasis drives ferroptosis represents an urgent scientific imperative[13].

Recent research indicates that the targeted induction of pathological calcium overload within cells can significantly promote immunogenic cell death (ICD) in TNBC[14]. Among the numerous intervention agents capable of disrupting intracellular calcium homeostasis, Nitric oxide(NO) is a potent redox signaling molecule capable of disrupting this Ca^2^ homeostasis[15]. Elevated NO levels drive tumor cell death by inducing concurrent oxidative stress and cytoplasmic Ca^2^ overload[11,16]. Thus, NO holds immense potential for metabolic intervention. Conventional NO donors, however, suffer from non-specific diffusion, preventing the precise targeting of subcellular microenvironments. An ideal NO doner drug requires a dual mechanism. It must release abundant cytoplasmic NO to trigger widespread Ca^2^ overload and simultaneously localize to specific organelles to disrupt confined Ca^2^□ stores. This synergistic approach would generate catastrophic intracellular oxidative stress, ultimately driving the cell into ferroptosis.

To address this issue, we synthesized 3A72, a novel furoxan-coumarin based NO donor. Here we investigate its pharmacological efficacy against TNBC cell proliferation. Furthermore, our study systematically elucidates the mechanisms by which 3A72 induces ferroptosis, including the disruption of calcium homeostasis at MAMs and interference with phospholipid metabolism. This provides experimental evidence for the development of next-generation anti-tumor drugs that target organelle crosstalk.

## 2 Materials and methods

### 2.1. Cell lines and antibodies

The triple-negative breast cancer cell lines 4T1 and MDA-MB-468 (purchased from the Cell Bank of the Shanghai Branch of the Chinese Academy of Sciences) were cultured separately in RPMI-1640 complete medium (Service Biotechnology, Wuhan, China)containing 10% fetal bovine serum (Capricorn Scientific, Germany), 1% penicillin–streptomycin (P/S, Meilunbio, China), and 1 mM sodium pyruvate, and Dulbecco’s Modified Eagle Medium (DMEM, Service Biotechnology,Wuhan,China) containing 10% FBS and 1% P/S respectively. All cells were maintained in a constant-temperature incubator at 37 °C, 5% CO_2_ and saturated humidity. Cells were routinely passaged every 2–3 days; following trypsinization and collection of the pellet by centrifugation at 1,000 rpm for 3 minutes, they were seeded into new culture flasks at a 1:3 ratio for continued culture. All *in vitro* experiments were conducted using cells of generations 3 to 10. Information on the primary antibodies used in this study and their sources is as follows: σ R (cat. No.: A5479); GRP75 (cat. No.: A11256) and ACSL4 (cat. No.: A20414) was purchased from Abclonal. LPCAT3 (cat. No.:HA723171), IP_3_R (HA601456), and GPX4 (cat. No.:ET1706-45) antibodies were all purchased from Hua’an Biotechnology. The internal control β-actin (cat. No: GB11001-100) and GAPDH (cat. No: GB11002-100) antibodies were both purchased from Servicebio.

### 2.2. Cell Viability Assay

Seed cells at a density of 1×10^5^ cells/mL into 96-well plates and allow them to adhere overnight. For cell viability assays, add 3A72 at different concentration gradients and incubate for 24 hours. For rescue experiments, first add a specified concentration of inhibitor pretreated for 1 hour, such as Carboxy-PTIO (2-(4-carboxyphenyl)-4,4,5,5-tetramethylimidazoline-1-oxyl-3-oxide, Carboxy-PTIO, an NO scavenger), NAC (N-Acetyl-L-cysteine, an ROS scavenger, Beyotime, S0077), Fer-1 (Ferrostatin-1, an ferroptosis inhibitor, Beyotime, Y240805), and 2-APB (2-Aminoethoxydiphenyl borate, an inositol 1,4,5-trisphosphate receptor (IP_3_R) inhibitor, MCE, CAS No.: 524-95-8), and pre-treat for half an hour, followed by the addition of 3A72 for a 12-hour co-incubation. Upon completion of the drug treatment, the culture medium was discarded, and 150 µL of MTT solution (5 g/L) was added to each well, which was then incubated in the dark for 4 hours. Finally, the supernatant was aspirated, and 150µL of DMSO was added to each well to thoroughly dissolve the formazan crystals. The absorbance of each well was measured at a wavelength of 570 nm using a microplate reader to assess cell viability. The cell viability was determined by: Cell viability = (sample O.D.-blank O.D.)/ (control O.D.-blank O.D.).

Based on the IC_50_ values determined via the MTT viability assay, subsequent mechanistic investigations were performed using concentrations of 5× and 10× IC50 of Compound 3A72. This treatment was adopted to evaluate early pharmacological events and mechanisms of action, thereby minimizing interference from late-stage cell death. Treatment durations (e.g., 2 h or 4 h) were selected based on the kinetic characteristics of each cellular event, as detailed in the respective methodologies.

### 2.3. Colony formation assay

Cells in the logarithmic growth phase were seeded at a density of 5,000 cells per well in a six-well plate. After attachment overnight, the cells were treated with 3A72 drug-containing medium at the specified concentration for 48 hours. Culturing was then continued in drug-free complete medium for 10–14 days (with medium changes every 3 days) until visually detectable cell colonies formed. After rinsing with PBS, the plates were fixed with 4% paraformaldehyde (Servicebio, G1101) for 20 minutes and stained with and stained with 0.1 % crystal violet (Beyotime, C0121) for 10 min.

Finally, the samples were rinsed with deionized water, air-dried, and imaged. Valid colonies comprising 50 or more cells were counted, and the colony formation rate was calculated as ‘(number of colonies / number of seeded cells) × 100%’ to assess the effect of 3A72 on long-term cell proliferation and survival.

### 2.4. EdU-DNA Synthesis Experiment

Cells in the logarithmic growth phase were seeded into a six-well plate at a density of 1×10^5^ cells per well. After the cells had attached and undergone the appropriate drug treatment, EdU working solution was added, and the plates were incubated for 2 hours. Following washing, the plates were fixed with 4% paraformaldehyde at room temperature for 20 minutes, followed by permeabilization with 0.3% Triton X-100 (Beyotime, P0096) for 15 minutes. Subsequently, add the Click Additive reaction solution containing Azide 488 fluorescent dye (CellorLab, Shanghai, China) and incubate at room temperature in the dark for 30 minutes. After washing with PBS containing 3% BSA to remove unbound dye, add Hoechst 33342 (Beyotime, C1022) stain and incubate at room temperature in the dark for 10 minutes to counterstain the cell nuclei. Finally, after thorough washing with PBS, images were acquired using an inverted fluorescence microscope (Olympus, Japan). Newly synthesized DNA appears as green fluorescence, while cell nuclei appear as blue fluorescence. The levels of DNA synthesis and proliferative activity in each group were quantitatively assessed by calculating the proportion of EdU-positive cells relative to the total cell population.

### 2.5. Determination of intracellular NO levels

Take 1×10^5^ cells per well from a culture in the logarithmic growth phase and seed them into a six-well plate. Once the cells have completely attached to the surface, carry out the corresponding drug treatments in accordance with the experimental design. Subsequently, dilute the DAF-FM DA probe (Beyotime, S0019S) solution to a working concentration of 1:1000 using serum-free medium, then add the diluted probe working solution to the well plate and incubate in a cell culture incubator at 37°C in the dark for 1 hour to allow the probe to fully enter the cells and bind to intracellular nitric oxide. Upon completion of the incubation, aspirate the probe-containing working solution and wash the cells three times with pre-chilled PBS to remove unbound free probe. Hoechst 33342 stain solution(Beyotime, C1022) is then added to count-stain the cell nuclei; after incubating for 10 minutes at room temperature in the dark, the cells are washed three more times with PBS. Finally, they are observed under an inverted fluorescence microscope, where total intracellular NO appears as green monomers and the cell nuclei as blue monomers.

### 2.6. Determination of mitochondrial superoxide content

Cells in the logarithmic growth phase were seeded into a six-well plate at a density of 2×10^5^ cells per well. After overnight attachment, the cells were treated with drugs at specified concentration gradients for 1 hour. After washing with PBS, add the MitoSO™ Red probe (diluted 1:1500, Beyotime, S0061S) followed by the Mito-Tracker Green probe (diluted 1:2000, Beyotime, C1048) and incubate for 30 minutes and 20 minutes respectively. After thoroughly washing the cells in each group with PBS to remove unbound probe, images were immediately captured using an inverted fluorescence microscope (Olympus, Japan), mitochondria appear as green fluorescence, while superoxide within the mitochondria appears as red fluorescence. Image analysis software was then used to quantitatively determine the average fluorescence intensity, thereby assessing the relative content of superoxide within the mitochondrial matrix.

### 2.7. Determination of intracellular ROS levels

Cells in the logarithmic growth phase were seeded at a density of 2×10^5^ cells per well in a six-well plate. After overnight confluence, the original culture medium was discarded, and the cells were incubated for 2 hours in drug-containing medium at a series of concentration gradients. Upon completion of the incubation, the original medium was aspirated, and the cells were gently washed three times with PBS pre-warmed to 37 °C to remove impurities from the residual medium. Following the instructions for the DCFH-DA( Beyotime, S1105S) fluorescent probe kit for detecting reactive oxygen species(ROS), dilute the solution 1:1000 in phenol red-free medium, add it to the six-well plates and incubate in the incubator for 20 minutes; Subsequently, the dye was discarded, and the cells were washed three times with PBS.

Hoechst 33342 (Beyotime, C1022) stain was then added to counterstain the cell nuclei; after incubation for 10 minutes at room temperature in the dark, the cells were washed three more times with PBS. Finally, the samples were observed under an inverted fluorescence microscope (Olympus, Japan), where total intracellular ROS appeared as green monomers and the cell nuclei as blue monomers.

### 2.8. Western blot experiment

Total proteins from cells or tissues were extracted using RIPA lysis buffer containing protease inhibitors, and their concentrations were standardized using the BCA method. Equal volumes of protein samples were separated by 10% SDS-PAGE electrophoresis(Beyotime, P0014A) and electro transferred onto a PVDF membrane(Beyotime, P0965). After blocking with skimmed milk at room temperature for 2 hours, a specific primary antibody was added and incubated overnight at 4°C. Following washing of the membrane with TBST(Beyotime, ST673), the corresponding HRP-labelled secondary antibody was added and incubated at room temperature for 1.5 hours. Finally, ECL chemiluminescent reagent(Meilunbio, MA0186) was added dropwise to develop the signal, and the grey-scale values of the bands were acquired and analyzed using Image Lab software. GAPDH and β-actin were used as internal controls to calculate the relative expression levels of the target proteins.

### 2.9. C11-BODIPY Staining Detection

After adding 3A72 to cells seeded in a six-well plate and left to culture overnight, aspirate the old culture medium and wash three times with PBS. Add a 1:1000 dilution of the C11-BODIPY probe (BDPY 581/591 C11, Beyotime, S0043S) working solution to each well, then place the plate in a 37°C cell culture incubator and incubate in the dark for 30 minutes. Subsequently, aspirate the probe working solution and wash with PBS to remove any free probe that has not been fully taken up by the cells. Add serum-free medium or PBS to cover the cells, and immediately capture images under a fluorescence microscope or confocal microscope. Under red light, unoxidized lipids are excited and under the green light channel, oxidized lipids are visualized.

### 2.10. Determination of Intracellular GSH and MDA Levels

Collect cells from each group and lyse them; after centrifugation, collect the supernatant and determine the total protein concentration using the BCA method. The determination of intracellular glutathione (GSH) and malondialdehyde (MDA) levels was carried out strictly in accordance with the instructions for the Beyotime assay kit. GSH was measured by GSH and GSSG detection kits(Beyotime, S0053) using the DTNB method: the deproteinized supernatant was mixed with the working solution, allowed to react at room temperature for 5 minutes, followed by the addition of NADPH, and the absorbance was measured at 412 nm using a microplate reader. The MDA assay is measured by Lipid Oxidation (MDA) Detection kit (Beyotime, S0131S) employed the TBA method: the supernatant was thoroughly mixed with the TBA working solution, heated at 100 °C for 100 minutes, cooled and centrifuged, and the absorbance measured at 532 nm. The concentrations of the final products were normalized according to the total protein concentration of each sample.

### 2.11. JC-1 Staining Assay

Changes in mitochondrial membrane potential were assessed using the JC-1 Fluorescent Probe Kit (Beyotime, C1475S). Cells seeded overnight in six-well plates were treated with 3A72 for 1 hour; the old culture medium was then aspirated, and the cells were washed three times with PBS. A prepared JC-1 staining working solution was added, and the plates were incubated at 37 °C in the dark for 20 minutes. Upon completion of the incubation, the staining solution was aspirated, and the cells were washed three times with pre-chilled PBS buffer; images were then immediately captured from randomly selected fields of view under an inverted fluorescence microscope (Olympus, Japan). Within undamaged mitochondria, JC-1 forms aggregates and emits red fluorescence; when the membrane potential is depolarized due to damage, JC-1 exists as monomers and emits green fluorescence. Finally, the average fluorescence intensity of the images from each group was analyzed using ImageJ software, and the extent of mitochondrial damage was quantitatively assessed by calculating the relative ratio of red to green fluorescence intensities.

### 2.12. Ferric Ion RhoNox-1 Probe Assay

The RhoNox-1 ferric ion fluorescent probe (MCE, CAS No.: 1447815-38-4) was used to assess the level of intracellular free ferrous ions (Fe^2^□) accumulation. Cells seeded in six-well plates and left to grow overnight were treated with 3A72 for 2 hours, after which the old culture medium was aspirated and the cells were washed three times with PBS. Subsequently, an appropriately diluted RhoNox-1 staining solution was added, and the cells were incubated in the dark at 37°C in a cell culture incubator for 60 minutes. Upon completion of the incubation, the staining solution was aspirated, and the cells were washed twice with PBS to thoroughly remove any free probe. Next, a diluted Hoechst 33342(Beyotime, C1022)staining solution was added to counterstain the cell nuclei; after incubation for 10 minutes at room temperature in the dark, the cells were washed three times with PBS. The samples were then immediately examined under an inverted fluorescence microscope, and images were captured from randomly selected fields of view. Upon reaction with Fe^2^□ specifically accumulated within the cells, the RhoNox-1 probe emits a bright orange-red fluorescence. Finally, ImageJ software was used to analyze the mean fluorescence intensity of the images from each group to quantitatively assess the relative levels of ferrous ions within the cells.

### 2.13. Subcutaneous Xenograft Tumors in Vivo

To establish a subcutaneous xenograft tumor model of triple-negative breast cancer, we collected 4T1 cells in the logarithmic growth phase, washed them, and resuspended them in PBS on an ice bath. Immediately thereafter, 1 × 10^6^ of 4T1 cells per milliliter were subcutaneously injected into the right dorsal region of 4–6-week-old female BALB/c mice that had been shaved, with 100 µL injected per mouse. One week after tumor implantation, tumor dimensions were measured periodically using a vernier caliper, and tumor volume was calculated using the formula V = (L × W^2^) / 2, where L and W represent the longest and shortest diameters of the tumor, respectively. For the *in vivo* efficacy evaluation, mice were randomized and treated with Compound 3A72 at a low dose (3 mg/kg) and a high dose (6 mg/kg), alongside a vehicle control and a positive control (Paclitaxel, PTX). This concentration gradient was strategically selected based on preliminary dose-tolerance tests to establish the dose-response relationship and delineate the optimal therapeutic window, balancing maximum anti-tumor efficacy with tolerable systemic toxicity. When the mean tumor volume reached 80–100 mm³, the tumor-bearing mice were randomly divided into a control group, two 3A72 treatment groups (3mg/kg and 6mg/kg), and a paclitaxel group (10mg/kg) for subsequent drug administration, with five mice in each group.

The experiment was terminated once a significant difference in tumor volume was observed. At the end of the experiment, the mice were euthanized; the tumors and major organ(heart, liver, spleen, lung, kidney) were rapidly excised and either frozen or fixed for subsequent histopathological assessment and analysis of the underlying mechanisms.

### 2.14. Bioinformatics and Gene Co-expression Analysis

Gene expression profiles and clinical data for breast invasive carcinoma (BRCA) were obtained from The Cancer Genome Atlas (TCGA) dataset. The UALCAN interactive web portal[17,18] was utilized to evaluate the differential mRNA expression levels and prognostic significance of the target genes in BRCA patients. To investigate functional co-expression, the LinkedOmics database[19] was employed to analyze the Pearson correlation between *ITPR3* (which encodes IP_3_R) and *ACSL4* expression in the TCGA-BRCA cohort. Furthermore, a protein-protein interaction (PPI) network was constructed using the STRING database[20], followed by Gene Ontology (GO) functional enrichment analysis to elucidate the underlying biological processes and pathways associated with the targets.

### 2.15. Computational Molecular Docking

To predict binding affinities and interaction modes, molecular docking was systematically performed. Protein-ligand blind docking between Compound 3A72 and target proteins was conducted using the CB-Dock2 web server [21,22], which integrates cavity detection, docking, and homologous template fitting. For protein-protein interaction predictions, rigid-body docking between the ACSL4 and ITPR3 proteins was carried out using the ClusPro web server [23]. The docking poses were evaluated based on energy scores, and the optimal binding conformations were selected for further analysis. All three-dimensional (3D) molecular interactions and docking interfaces were visualized and rendered using UCSF ChimeraX software.

### 2.16. Statistical Analysis of Biological Data

All quantitative experimental data are expressed as mean ± standard deviation (Mean ± SD). GraphPad Prism software was used for statistical analysis and graphing of the data. Differences between two groups were compared using the unpaired Student’s t-test; comparisons between multiple groups were performed using one-way analysis of variance (ANOVA), combined with Tukey’s (or Dunnett’s) post-hoc tests. All *in vitro* cellular experiments were repeated independently at least three times. P < 0.05 was considered statistically significant (where *P < 0.05, P < 0.01, ***P < 0.001).

## 3 Experiment Results

### 3.1. 3A72 Inhibits the In Vitro Proliferation of TNBC Cells

This study aimed to evaluate the anti-tumor potential of the novel furazan-coumarin conjugate 3A72 (Fig. 1A). Firstly, the cytotoxic effect of 3A72 on two TNBC cell lines (MDA-MB-468 and 4T1) was assessed using the MTT assay, following 24-hour treatment with 3A72 at a range of concentrations, the compound was found to significantly inhibit cell viability, with the inhibitory effect increasing with rising concentration. IC_50_ values were calculated from the concentration-response curves; for both MDA-MB-468 (IC_50_ =17.48 nM) and 4T1 (IC_50_ = 121.8 nM), 3A72 was shown to significantly inhibit the proliferation of TNBC cells (Fig. 1B).

**Fig. 1.**
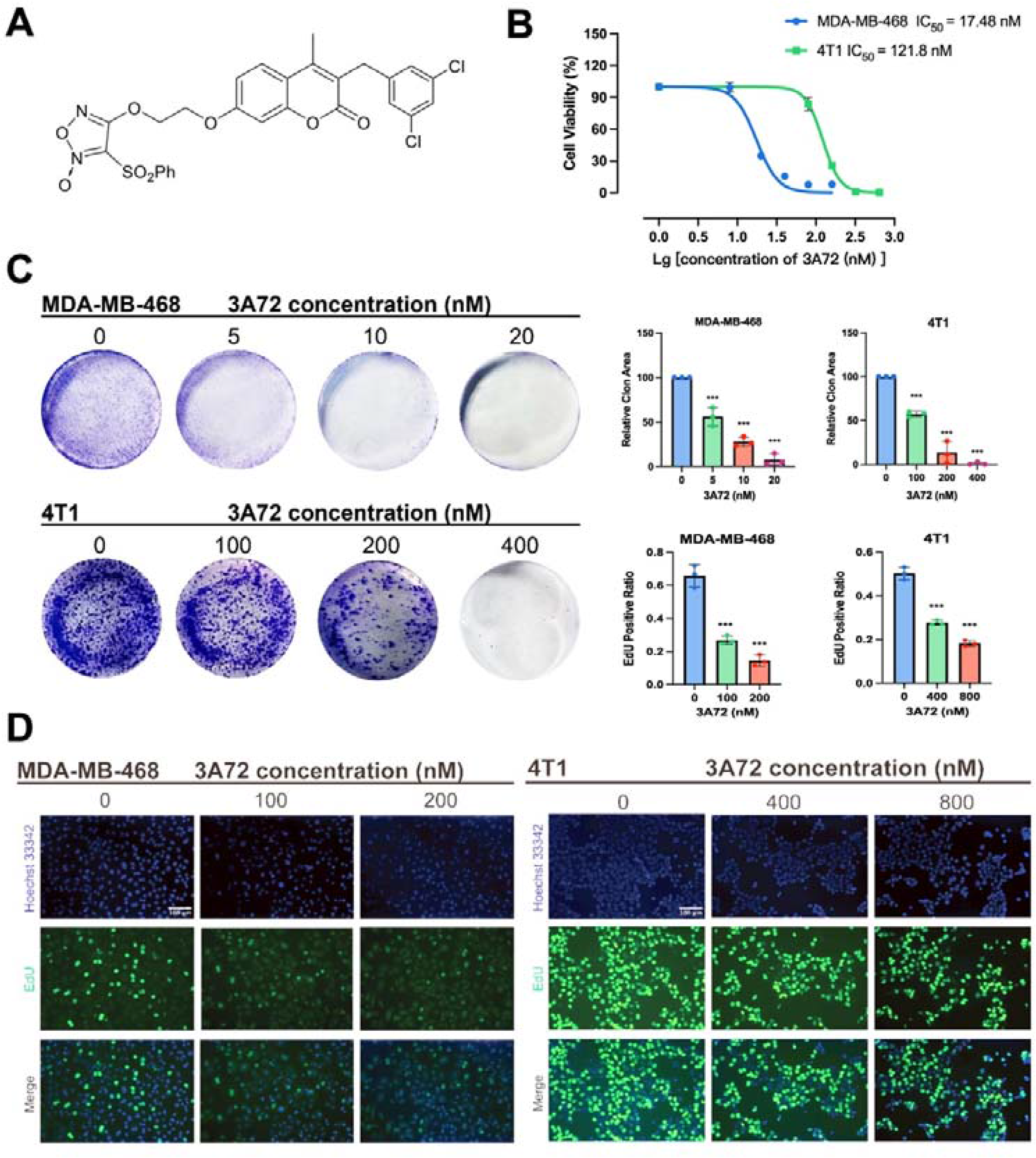
3A72 inhibits the proliferation of TNBC cells. (A) Chemical structure of 3A72. (B) Cell viability and IC values of MDA-MB-468 and 4T1 cells treated with various concentrations of 3A72. (C) Representative images and quantitative analysis of colony formation assays in MDA-MB-468 cells and 4T1. (D) Representative fluorescence images and quantification of EdU incorporation assays in MDA-MB-468 and 4T1 cells. Data are expressed as mean ± SD (n = 3). *p < 0.05, **p < 0.01, ***p < 0.001 vs. the control group.

To evaluate the long-term effects of 3A72 on TNBC cell proliferation, we conducted colony formation assays. The results showed that 3A72 inhibited colony formation in MDA-MB-468 and 4T1 cells in a dose-dependent manner (Fig. 1C). Furthermore, EdU incorporation assays revealed that 3A72 significantly inhibited DNA synthesis in both cell lines, ultimately leading to TNBC cell death (Fig. 1D). These results indicate that compound 3A72 effectively inhibits the proliferation and clonogenic capacity of TNBC cells *in vitro* and interferes with their DNA synthesis.

### 3.2. 3A72 causes cytoplasmic calcium overload via the release of NO

Based on its furazan-coumarin chemical structural core, 3A72 is considered to be a novel NO donor. Accordingly, we hypothesized that its role in inducing cell death in TNBC cells might be related to the release of NO. We first utilized the DAF-FM DA fluorescent probe to detect changes in intracellular total NO levels. The results showed that, following treatment with 3A72, the mean fluorescence intensity (MFI) of total NO in MDA-MB-468 and 4T1 cells increased significantly in a concentration-dependent manner (Fig. 2A).

**Fig. 2.**
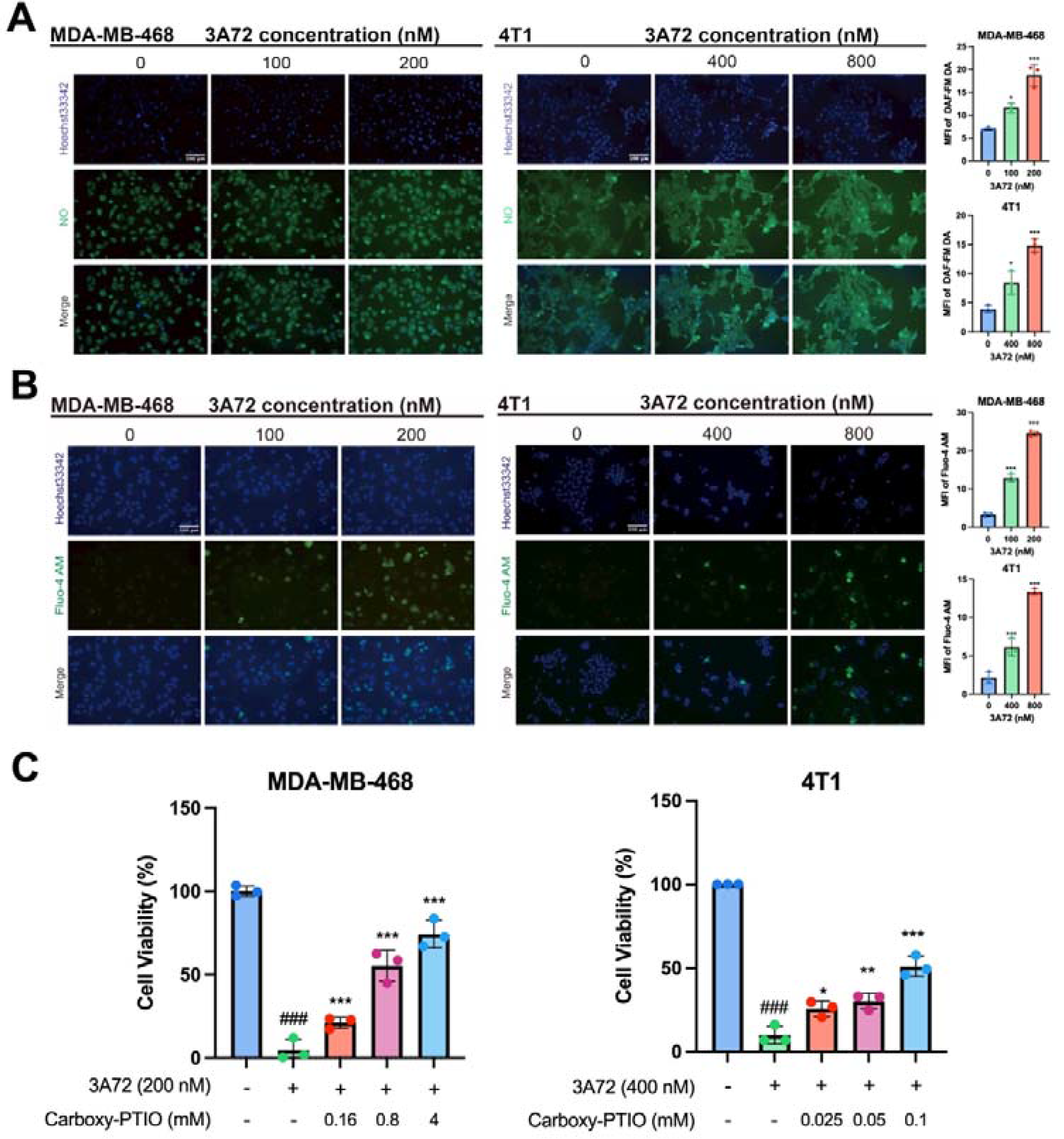
3A72 induces cytosolic calcium overload via NO release. (A) Intracellular NO levels in MDA-MB-468 and 4T1 cells treated with 3A72 detected by DAF-FM DA staining. (B) Cytosolic calcium levels in 3A72-treated cells measured by Fluo-4 AM staining. (C) Viability of cells pre-treated with NO scavenger Carboxy-PTIO and then co-treated with 3A72. Data are presented as mean ± SD (n = 3). ^###^p < 0.001 vs. control; * p < 0.05, ** p < 0.01, *** p < 0.001 vs. 3A72-treated group. MFI: Mean fluorescence intensity.

Studies have shown that NO promotes endoplasmic reticulum calcium release via S-nitrosylation of RyR receptors, and in concert with downregulation of plasma membrane calcium ATPase (PMCA) expression, blocks calcium efflux, thereby inducing intracellular calcium overload[24]. In our experiment, we observed via Fluo-4AM fluorescence imaging that cytoplasmic calcium ion concentrations increased significantly following 3A72 treatment (Fig. 2B). However, the rescue experiment using the NO scavenger Carboxy-PTIO revealed that while Carboxy-PTIO significantly improved the survival rate of TNBC cells, it failed to fully restore it, indicating that 3A72-induced cell death depends not only on the direct action of NO but also on other key downstream signaling pathways (Fig. 2C).

In summary, 3A72 mediates cytoplasmic calcium overload via the release of NO, thereby inducing TNBC cell death. However, given that NO scavengers failed to completely reverse cell viability, this suggests that the lethal effects of 3A72 may also involve the synergistic participation of other molecular mechanisms.

### 3.3. 3A72 acts on IP_3_R and ACSL4 to induce mitochondrial calcium overload by regulating the function of the MAMs calcium channel

Among the treatment regimens targeting the heterogeneity of TNBC, MAMs have attracted significant attention as key structures involved in calcium homeostasis and metabolic regulation[11,25]. Using the UALCAN online database, we analyzed the expression of *ITPR3* (which encodes the IP_3_R protein) and *ACSL4* across different molecular subtypes of breast cancer in the TCGA-BRCA cohort. We found that, compared with other breast cancer subtypes, *ITPR3* and *ACSL4* were significantly over-expressed in TNBC tissues comparing to other subtypes (Fig. 3A–B), suggesting that these two genes hold significant clinical value in the pathology of TNBC. From a biological perspective, IP_3_R, as a key component of the MAMs calcium transport axis, directly mediates the transport of calcium ions from the endoplasmic reticulum to the mitochondria and acts as a central regulator of cellular calcium homeostasis; ACSL4, as a key enzyme in lipid metabolism, is highly associated with MAMs function[26] by remodeling the phospholipid microenvironment in the MAMs region[27], thereby providing essential support for the assembly and structural stability of this calcium transport complex. Given their pivotal roles in maintaining the function of the MAMs calcium axis, we hypothesize that they represent key downstream targets for the anti-tumor activity of 3A72.

**Fig. 3.**
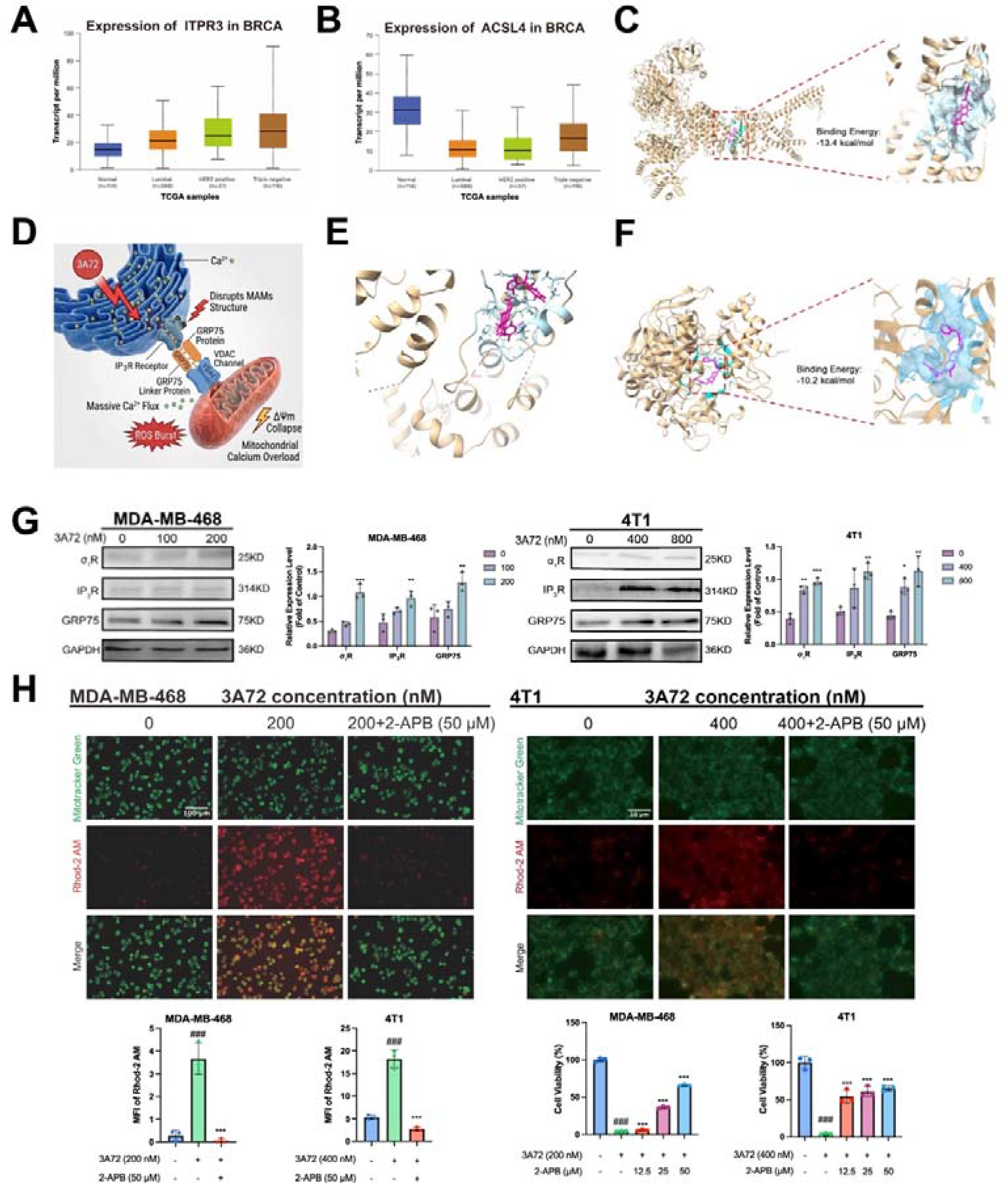
3A72 induces mitochondrial calcium overload by targeting IP3R and ACSL4 at MAMs. (A, B) Expression of ITPR3(IP3R) (A) and ACSL4 (B) in breast cancer subclasses based on TCGA data. (C, E, F) Molecular docking of 3A72 with its putative targets. (D) Schematic illustration of 3A72-induced disruption of MAMs and mitochondrial calcium overload. (G) Protein levels of MAMs-associated proteins (σ1R, IP3R, and GRP75) in 3A72-treated cells. (H) Mitochondrial calcium levels measured by Rhod-2 AM and MitoTracker Green co-staining. (I) Viability of cells co-treated with 3A72 and the IP3R inhibitor 2-APB. Data are presented as mean ± SD (n = 3). ###p < 0.001 vs. control; *p < 0.05, ***p < 0.001 vs. 3A72-treated group. MFI: Mean fluorescence intensity.

To further investigate the specific core targets and molecular mechanisms by which 3A72 and the NO it releases exert their anti-tumor effects, we utilized molecular docking techniques to simulate the binding patterns between 3A72 and the potential core target proteins IP_3_R and ACSL4. The results showed that 3A72 is able to precisely anchor itself and enter the active binding pocket of these target proteins; the binding energy with the IP_3_R protein was as low as −13.4 kcal/mol, while that with the ACSL4 protein was −10.2 kcal/mol, both of which were lower than −8 kcal/mol (Fig. 3C, F), indicating extremely strong binding affinity between them. Further 3D conformation and amino acid interaction analyses between 3A72 and IP_3_R (Fig. 3E) revealed that the 3A72 molecule forms a tight network of hydrogen and hydrophobic interactions with several key cysteine residues within the pocket; this not only maintains the structural stability of the complex but also strongly suggests that IP_3_R and ACSL4 are likely the key direct sites through which 3A72 exerts its pharmacological effects.

To investigate the impact of 3A72’s interactions with IP_3_R and ACSL4 on calcium signaling, we assessed changes in MAMs structural proteins and mitochondrial calcium levels. Western blot analysis confirmed that 3A72 significantly increased the expression of key MAMs calcium channel proteins, such as σ R, IP_3_R, and GRP75 (Fig. 3G), suggesting that 3A72 disrupts the structural integrity of the contact interface between the endoplasmic reticulum and mitochondria. Further analysis using a mitochondria-specific calcium probe revealed that, following treatment with 3A72, the fluorescence intensity of calcium ions within the mitochondrial matrix was significantly enhanced, confirming that, following 3A72 treatment, calcium ions underwent a massive abnormal transmembrane influx into the mitochondria from the endoplasmic reticulum via activated MAMs, leading to mitochondrial calcium overload. Furthermore, pre-treatment of cells with the IP R inhibitor 2-APB prior to 3A72 administration significantly reversed the activation of the MAM calcium channel and the 3A72-induced calcium overload (Fig. 3H). Furthermore, pre-treatment of TNBC cells with 2-APB at a concentration of 50 µM prior to co-incubation with 3A72 showed that 2-APB significantly reversed the cytotoxic effects of 3A72 (Fig. 3I). In summary, 3A72 binds to key MAM proteins and excessively activates the IP_3_R-mediated calcium transport axis, leading to severe mitochondrial calcium overload, which is the key mechanism triggering subsequent cell death.

### 3.4. 3A72 activates IP_3_R to mediate a calcium-ROS cascade, triggering mitochondrial dysfunction and oxidative stress

Mitochondria are the key organelles responsible for regulating redox homeostasis and energy metabolism within cells. Physiological calcium concentrations are fundamental to maintaining their normal function, and therefore a sustained increase in intracellular calcium ion concentrations directly disrupts mitochondrial oxidative balance[28]. Studies have shown that mitochondrial calcium overload triggers a burst of ROS by activating the dehydrogenase activity in the tricarboxylic acid (TCA) cycle, causing the electron transport chain (ETC) to operate at an excessive rate and leading to electron leakage[29]. Previous findings indicate that 3A72 induces mitochondrial calcium overload by binding to the IP_3_R protein, thereby activating the MAMs structural calcium channel. We further investigated whether mitochondrial calcium overload leads to the production of large amounts of ROS by mitochondria.

Experimental results showed that following treatment with 3A72, mitochondrial ROS levels increased significantly, and the IP_3_R inhibitor 2-APB could markedly inhibit the 3A72-induced mitochondrial ROS overload (Fig. 4A), demonstrating that the generation of large amounts of ROS in mitochondria is caused by 3A72-mediated activation of IP_3_R. Concurrently, the large amount of ROS generated in the mitochondria rapidly triggered a decline in the mitochondrial membrane potential (MMP); JC-1 staining results clearly demonstrated the transition from red polymers, indicative of a high membrane potential, to green monomers, indicative of a low membrane potential (Fig. 4B). The ratio of red polymers to green monomers decreased as the concentration of 3A72 increased, confirming the rapid collapse of mitochondrial function under oxidative damage. As the mitochondrial membrane structure was damaged, large amounts of ROS accumulated within the mitochondria leaked into the cytoplasm, thereby triggering a more widespread accumulation of cytoplasmic ROS. DCFH-DA staining results indicate that, Following 3A72 treatment, ROS levels in the cytoplasm exhibited a significant dose-dependent increase (Fig. 4C); furthermore, pretreatment with the ROS scavenger NAC completely reversed the anti-proliferative effect of 3A72 on TNBC (Fig. 4D), demonstrating that following binding to IP_3_R, 3A72 activated MAMs-driven mitochondrial oxidative damage, which subsequently cascaded into the cytoplasm via mitochondrial ROS leakage, leading to ROS overload and thereby establishing the critical oxidative stress environment required for subsequent lipid peroxidation.

**Fig. 4.**
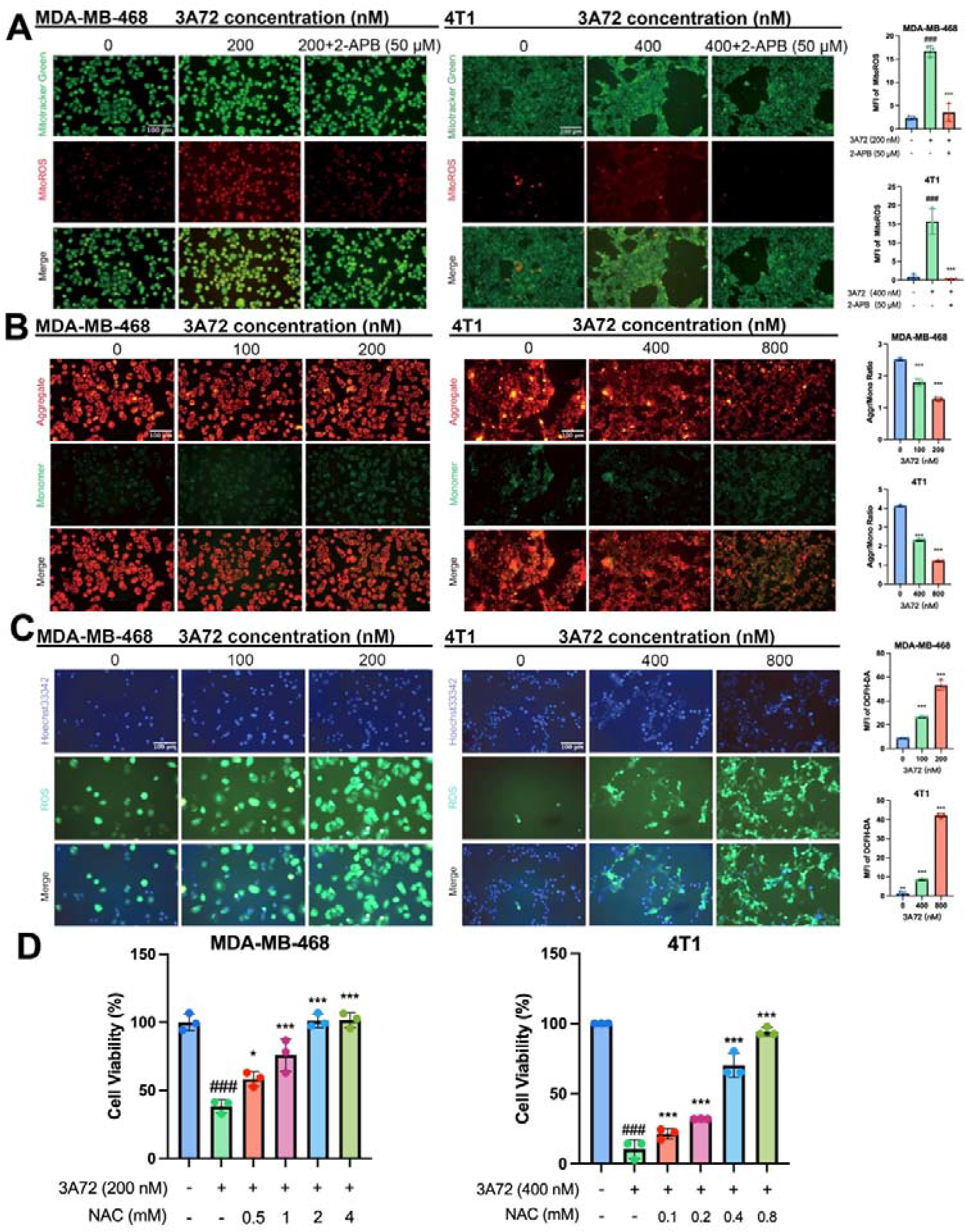
3A72 induces mitochondrial dysfunction and oxidative stress via the IP_3_R-mediated calcium-ROS cascade. (A) Mitochondrial ROS levels assessed by Mitotracker Green and MitoROS. (B) Mitochondrial membrane potential measured by JC-1 staining. (C) Intracellular ROS levels detected by DCFH-DA staining. (D) Cell viability after co-treatment with 3A72 and the ROS scavenger NAC. Data are mean ± SD (n = 3).^###^p < 0.001 vs. control; * p < 0.05, *** p < 0.001 vs. 3A72 group. MFI: Mean fluorescence intensity

### 3.5. 3A72 drives the phospholipid remodeling pathway to induce lipid peroxidation in TNBC cells

Based on the molecular docking results presented in 3.3, which demonstrated 3A72’s extremely high affinity for the ACSL4 and IP_3_R proteins, it is suggested that 3A72 may regulate the structural homeostasis of MAMs and lipid metabolism by interfering with the functions of IP_3_R and ACSL4. As a key enzyme regulating cell membrane lipid composition, ACSL4 is not only the rate-limiting step in lipid remodeling but also a core molecule determining the cell’s sensitivity to ferroptosis[30]. To investigate how 3A72 precisely regulates this metabolic node, we first analyzed the interaction network of ACSL4 using the STRING database (Fig. 5A). PPI network and GO functional enrichment analyses revealed that ACSL4 is highly enriched in the structural components of MAMs and exhibits strong physical interactions with key calcium channel protein’s gene such as *ITPR3* (Fig. 5B). This suggests that ACSL4 most likely maintains the microenvironmental homeostasis of MAMs by integrating calcium signaling with lipid metabolism. We further performed protein-protein docking at the three-dimensional atomic level to confirm that ACSL4 and IP_3_R can form a physically interactive complex with a highly stable conformation (lowest energy weighted score: −1026) (Fig. 5D), demonstrating the structural potential for the formation of a stable complex between the two and further supporting the suggestion of a physical link revealed in the aforementioned network analysis.

**Fig. 5.**
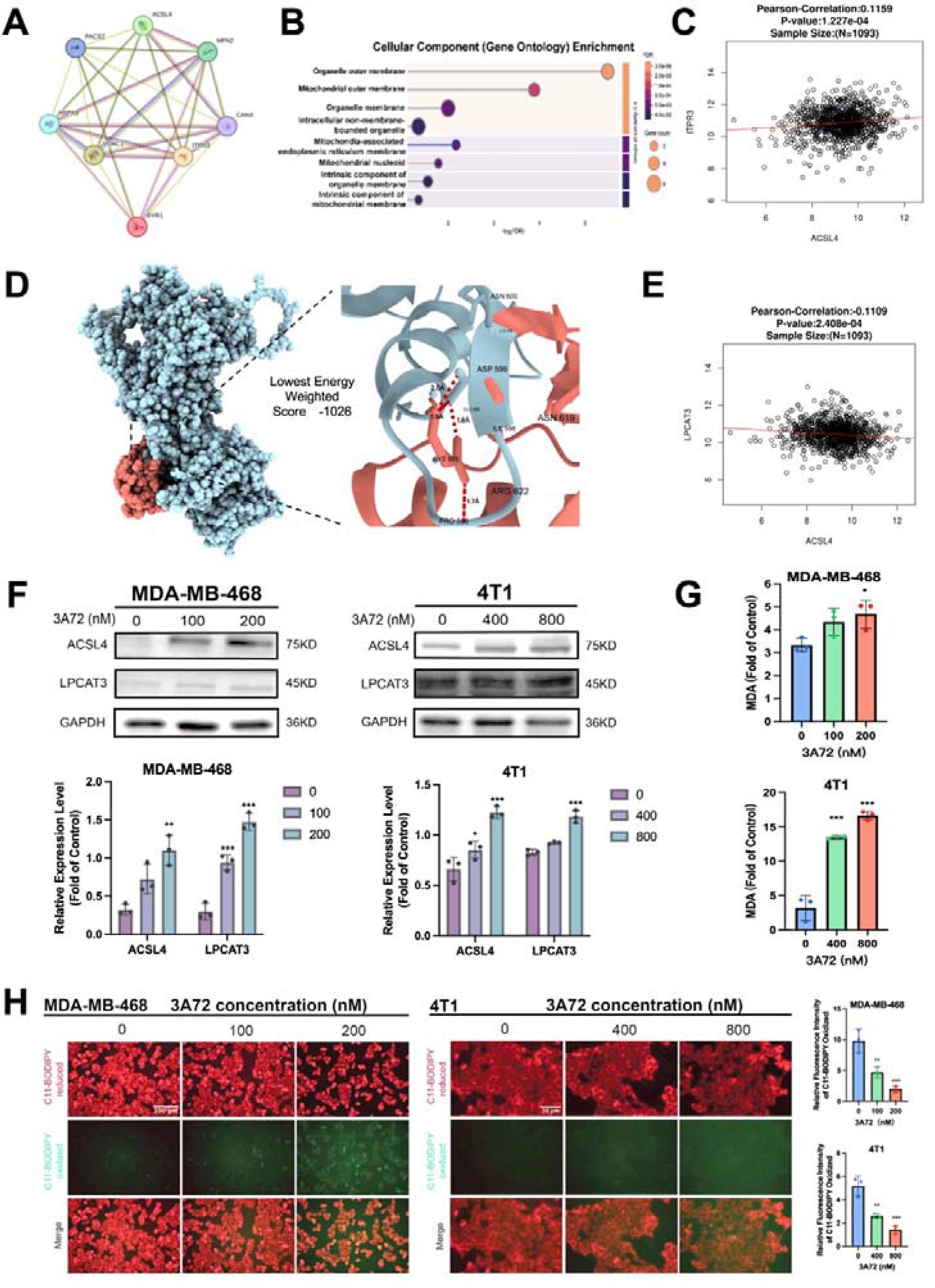
3A72 induces lipid peroxidation via phospholipid remodeling. (A) Protein-protein interaction network of ACSL4, ITPR3, and related interacting proteins (PACS2, MFN2, CANX, RYR1, VDAC1, and HSPA9). (B) Gene Ontology (GO) enrichment analysis of cellular components. (C, E) Correlation analysis of ACSL4 with ITPR3 (C) and LPCAT3 (E). (D) Molecular docking simulation between ACSL4 and IP3R. (F) Western blot analysis of ACSL4 and LPCAT3 protein levels in MDA-MB-468 and 4T1 cells. (G) Relative MDA levels in 3A72-treated cells. (H) Lipid peroxidation evaluated by C11-BODIPY staining. Data are presented as mean ± SD (n = 3). * p < 0.05, ** p < 0.01, *** p < 0.001 vs. control. MFI: Mean fluorescence intensity.

Based on this structural foundation, we further analyzed the expression profile of ACSL4 in clinical samples. Data from the TCGA-BRCA cohort revealed that, in TNBC tumors, ACSL4 exhibited a significant negative correlation with LPCAT3, a key enzyme downstream of lipid remodeling (Pearson r = −0.1109, P < 0.001), while showing a positive correlation with *ITPR3* (Pearson r = 0.1159, P < 0.001) (Fig. 5C, E). Clinical evidence suggests that, under physiological conditions, TNBC cells finely regulate ACSL4 and LPCAT3-mediated lipid metabolism to prevent excessive phospholipid remodeling from triggering lethal lipid peroxidation[30]. Experimental results indicate that treatment with 3A72 leads to a significant increase in the protein expression of ACSL4 and LPCAT3, demonstrating that 3A72, by activating the phospholipid remodeling process in lipid metabolism (Fig. 5F), causes an increase in the lipid metabolism intermediate MDA (Fig. 5G), thereby triggering intense lipid peroxidation in TNBC cells. We further confirmed through C11-BODIPY probe fluorescence imaging that, as the concentration of 3A72 increased, intracellular phospholipids underwent dramatic redox state changes, red fluorescence diminished, while green fluorescence increased significantly (Fig. 5H). In summary, 3A72 disrupts the lipid metabolic homeostasis of TNBC cells by activating the *ITPR3*– ACSL4 structural hub and by driving the ACSL4/LPCAT3-mediated phospholipid remodeling programmed, it ultimately induces irreversible lipid peroxidation.

### 3.6. 3A72 induces ferroptosis via lipid peroxidation

Lipid peroxidation is a key factor in triggering ferroptosis[31]; we therefore conducted further experimental studies on the induction of ferroptosis in TNBC cells by 3A72. Protein Western blot analysis was performed to detect changes in the expression of GPX4 and ACSL4, key regulatory proteins of ferroptosis. The results showed that, following treatment with 3A72, GPX4 levels in TNBC cells decreased significantly as the drug concentration increased, while ACSL4 levels rose significantly with increasing drug concentration, indicating that 3A72 activated the ferroptosis pathway (Fig. 6A); subsequently, using a ferrous ion-specific RhoNox-1 fluorescent probe, we found that treatment with 3A72 significantly increased ferrous ion levels within TNBC cells, whereas prior inhibition of the upstream IP_3_R/MAMs calcium pathway using 2-APB markedly reversed this accumulation of ferrous ions, confirming that iron ion accumulation depends on the activation of the upstream MAMs calcium pathway (Fig. 6B).

**Fig. 6.**
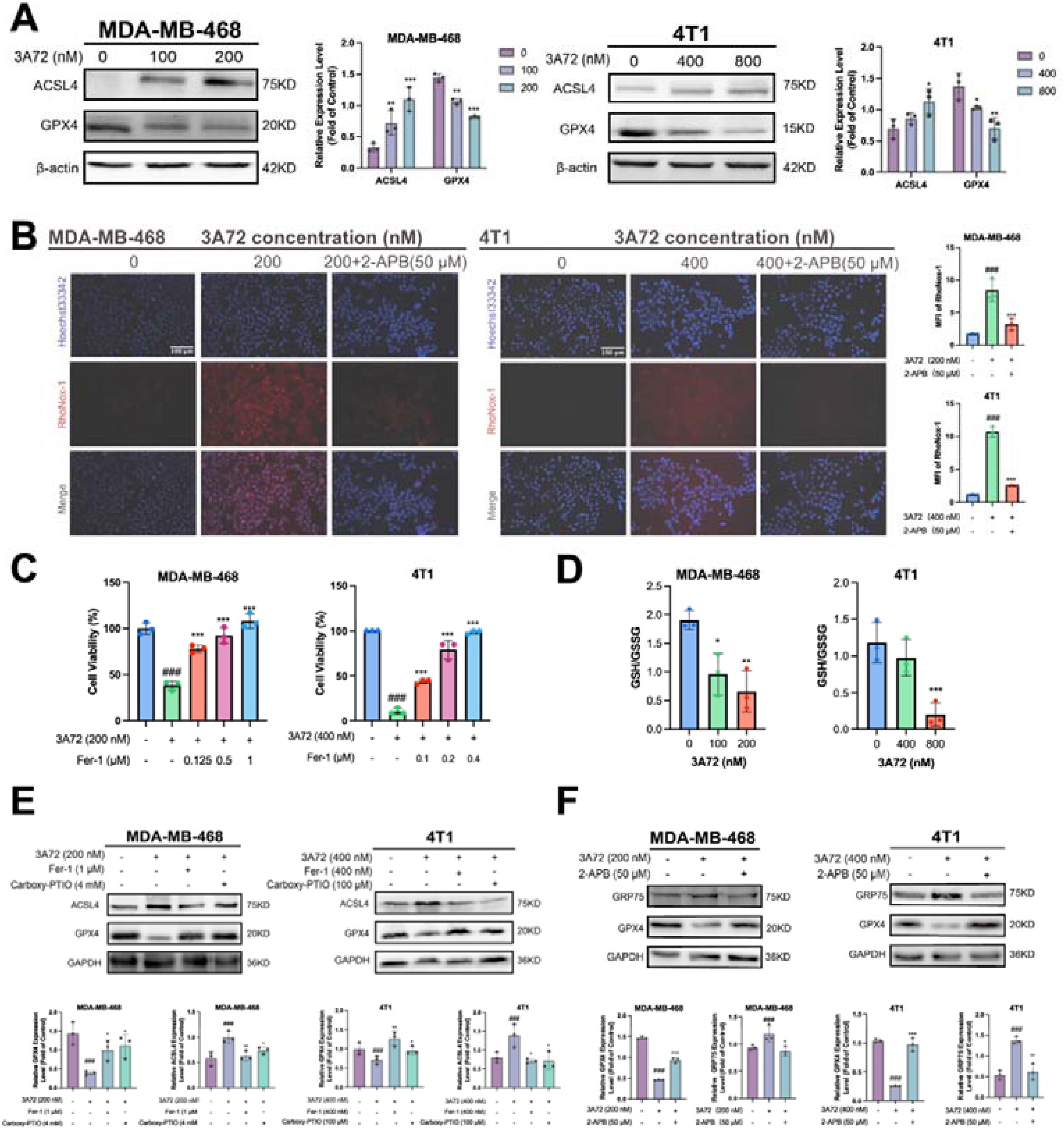
3A72 induces ferroptosis in TNBC cells. (A) Protein levels of ACSL4 and GPX4 in 3A72-treated MDA-MB-468 and 4T1 cells. (B) Intracellular Fe^2+^ levels detected by RhoNox-1 staining after 3A72 and 2-APB co-treatment. (C) Viability of cells co-treated with 3A72 and the ferroptosis inhibitor Fer-1. (D) GSH/GSSG ratio in 3A72-treated cells. (E,F) Western blot analysis of indicated proteins in cells co-treated with 3A72 and Fer-1, Carboxy-PTIO, or 2-APB. Data are presented as mean ± SD (n = 3). p < 0.05, ** p < 0.01, *** p < 0.001 vs. control or 3A72-treated group; ^###^p < 0.01, ^###^p <0.001 vs. control. MFI: Mean fluorescence intensity.

To further demonstrate that the inhibition of cellular activity was caused by the induction of ferroptosis, we pre-treated the cells with the ferroptosis-specific inhibitor Fer-1 before co-incubating them with 3A72; the results showed that inhibition of ferroptosis reversed the regulation of ferroptosis-related TNBC cell death caused by 3A72 (Fig. 6C). Analysis of the ratio of intracellular GSH to GSSG levels following treatment revealed that the GSH/GSSG ratio was significantly reduced in cells treated with 3A72, consistent with the characteristic depletion of glutathione and disruption of redox homeostasis during ferroptosis (Fig. 6D). These results indicate that 3A72 induces ferroptosis by triggering systemic oxidative stress within the cell, which in turn leads to lipid metabolism dysregulation. Furthermore, co-incubation of TNBC cells with Fer-1 or Carboxy-PTIO, respectively, in the presence of 3A72 markedly increased GPX4 protein expression while reducing ACSL4 expression (Fig. 6E), demonstrating that 3A72 activates a lipid peroxidation-mediated ferroptosis signaling pathway, leading to ferroptosis in TNBC cells. Pre-treatment of TNBC cells with 2-

APB followed by co-incubation with 3A72 similarly increased GPX4 protein expression and reduced GRP75 expression (Fig. 6F), demonstrating that IP_3_R-GRP75-mediated calcium overload within the MAMs leads to ferroptosis in TNBC cells.

### 3.7. Inhibition of TNBC proliferation by 3A72 in vivo

We further investigated the *in vivo* anticancer activity of 3A72 by establishing a 4T1 cell xenograft mouse model (Fig. 7A). The results indicated that 3A72 significantly restricted tumor growth in a dose-dependent manner. Experimental findings showed that, compared with the control group, the 3A72-treated groups (3 mg/kg and 6 mg/kg) not only significantly inhibited tumor proliferation but also achieved therapeutic effects equivalent to those of the classic first-line drug paclitaxel (administered twice weekly at 10 mg/kg) (Fig. 7B–D). At the same time, treatment-related adverse effects, such as weight loss, were less severe than in the paclitaxel group (Fig. 7E), with the animals in the latter group maintaining their body weight consistently near the normal baseline.

**Fig. 7.**
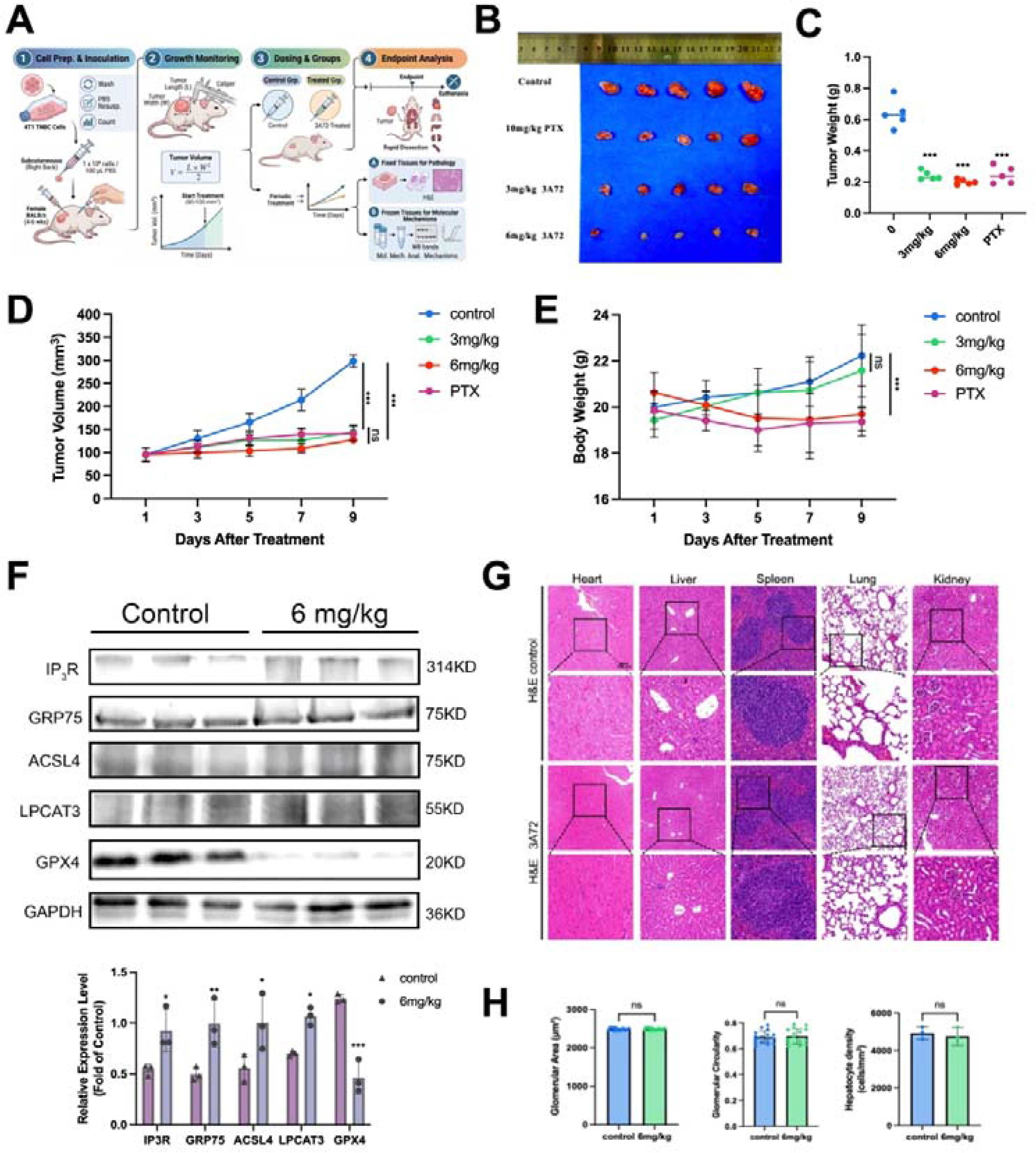
In vivo anti-tumor efficacy and safety evaluation of 3A72. (A) Schematic overview of the in vivo experimental design. (B, C) Representative images of excised tumors and their corresponding weights at the endpoint. (D) Tumor volume and (E) body weight growth curves of mice during the treatment period. (F) Western blot analysis and quantification of protein levels in tumor tissues. (G) Representative H&E staining images of major organs (heart, liver, spleen, lung, and kidney).(H) Quantitative analysis of glomerular area, glomerular circularity, and hepatocyte density. Data are presented as mean ± SD (n = 5). * p < 0.05, ** p < 0.01, *** p < 0.001 vs. control. ns: not significant.

Subsequently, based on the molecular mechanism by which 3A72 induces ferroptosis in TNBC via MAMs activation, as demonstrated in *in vitro* models, we also investigated the molecular mechanisms in mouse tumor tissues. Analysis via Western blotting confirmed that 3A72 intervention not only significantly suppressed the expression of classic tumor-inhibitory ferroptosis proteins GPX4, and upregulated the expression of ACSL4, but also markedly increased the expression of MAMs-related proteins (such as IP3R, GRP75, ACSL4) (Fig. 7F), indicating that regulating MAMs structure and cascading the initiation of ferroptosis constitutes the key molecular basis for 3A72’s inhibition of subcutaneous xenograft tumor growth.

To further evaluate the *in vivo* toxic side effects and safety profile of 3A72, tumor tissues and target organs (heart, liver, spleen, lung, and kidney) were collected at the end of the experiment, and histological sections were analyzed using hematoxylin and eosin (H&E) staining. As shown in Fig. 7G, cardiac muscle fibers in both the untreated group and the 3A72-treated group were tightly arranged; the structure of the hepatic lobules was clear and intact, with no obvious vacuolization, degeneration, or necrosis; the red and white pulp of the spleen were distinctly separated; the alveolar structure was morphologically normal, with no evidence of septal thickening or congestion; the morphology and structure of the glomeruli and renal tubules were both distinct, suggesting preliminarily that 3A72 did not induce significant systemic toxicity in mice; Quantitative histomorphometry analysis revealed that, compared with the control group, there were no significant differences in glomerular area, glomerular circularity, or hepatocyte density in the 6 mg/kg treatment group (Fig. 7H). These results indicate that treatment with a dose of 6 mg/kg did not cause any marked morphological changes or toxic damage to the microscopic tissue structures of the liver and kidneys.

## 4. Discussion

Although traditional chemotherapy has long been the primary treatment for TNBC, its efficacy is often compromised by drug resistance in patients with advanced-stage disease, which significantly reduces patients’ quality of life and survival rates. Consequently, the development of a safe anti-tumor drug based on non-conventional apoptotic pathways has become crucial for the treatment of TNBC. In this study, we designed and optimized a series of compounds based on the furan-coumarin core and found that one of them, compound 3A72, possesses a strong ability to inhibit the proliferation of TNBC cells both *in vitro* and *in vivo*, as well as to prevent normal DNA replication. Furthermore, 3A72 demonstrated a significant inhibitory effect on tumor proliferation in a mouse subcutaneous 4T1 xenograft model; it also showed good *in vivo* safety and the potential to induce ferroptosis in TNBC cells.

Recent studies have confirmed that high concentrations of NO can induce S-nitrosylation of RyR on the endoplasmic reticulum (ER) membrane, leading to excessive calcium release into the cytoplasm. This results in cytoplasmic calcium overload and inhibits calcium efflux, acting as a key driver of lethal cytoplasmic calcium overload[24]. Experimental results indicate that, under the influence of 3A72, the fluorescence intensity of NO within TNBC cells exhibits a significant concentration-dependent increase; concurrently, under the same conditions, this leads to an elevation in cytosolic calcium ion concentration, suggesting that this class of compounds may induce cell death by triggering cytosolic calcium overload via the release of NO. It is worth noting that the NO scavenger Carboxy-PTIO only partially reversed the 3A72-induced cytoplasmic calcium overload and cell death, demonstrating that, in addition to the crucial role of the NO release pathway, non-NO-dependent synergistic mechanisms also contribute to 3A72-induced TNBC cell death. Among the genes relevant to TNBC treatment, we found that the expression of IP_3_R and ACSL4 was significantly increased compared with other breast cancer subtypes. Furthermore, as the initiator protein in the structural calcium axis of MAMs, IP_3_R is highly associated with calcium transport [29]; simultaneously, the function of ACSL4 is highly correlated with MAMs[32]; the membrane phospholipid remodeling it mediates is not only a key driver of malignant metastasis in TNBC[33], but also a core lipid metabolism gene involved in the induction of ferroptosis. Given the important roles of IP_3_R and ACSL4 in regulating stress-induced cell death in TNBC, and to further investigate the direct molecular targets underlying 3A72-induced TNBC cell death, we conducted molecular docking simulations. The molecular docking results showed that 3A72 is able to precisely fit into the active site of the target proteins IP_3_R/ACSL4 with extremely high affinity, forming stable hydrogen bond interactions with key amino acid residues (such as Tyr815), with a binding energy to IP_3_R as high as −13.4 kcal/mol. This precise binding at the molecular level not only provides a solid structural basis for the delivery of NO by 3A72 but also directs its pharmacological action directly towards the microenvironment where the target protein is located.

Given that core target proteins such as IP3R are highly enriched in the mitochondria-associated endoplasmic reticulum membranes (MAMs) region within cells, upon further investigation into the specific subcellular mechanisms by which this class of compounds exerts its effects, we found that their function is closely linked to the homeostasis of the mitochondria-associated endoplasmic reticulum (MAMs). MAMs are specialized regions where material exchange takes place between the endoplasmic reticulum and mitochondria[34]. According to classical theory, calcium permeability at MAMs is strictly regulated by chaperones; that is, under resting conditions, σ R is typically anchored by inhibitory proteins such as the chaperone BiP[35], while IP_3_R at MAMs usually undergoes rapid metabolic turnover via the ubiquitin-proteasome system (UPS) to maintain low levels of basal expression[36]. In our experimental system, drug intervention with 3A72 not only significantly upregulated the overall expression of the stress response protein σ R but also further led to a surge in the protein levels of its downstream core target, IP_3_R. In summary, upon entry into TNBC cells, 3A72 binds to IP_3_R, promoting the release of σ R from inhibition and its enrichment in the MAMs region. This process blocks the ubiquitination-mediated degradation of IP_3_R and triggers the activation of the MAMs calcium channel, leading to sustained mitochondrial calcium overload[37]. The disruption of MAMs homeostasis caused by calcium imbalance constitutes the starting point of the molecular mechanism by which 3A72 specifically triggers TNBC cell death.

The core biochemical feature of ferroptosis activation is the massive accumulation of membrane lipid peroxidation, a process that must be driven synergistically by ACSL4 and LPCAT3. Phospholipid remodeling provides a large amount of substrate for lipid peroxidation by acting on specific polyunsaturated fatty acids (PUFAs)[26]. Based on omics-based cellular component enrichment analysis and sequence-based transmembrane topological structure prediction, we confirmed that ACSL4 possesses dual anchoring properties in both the endoplasmic reticulum and the outer mitochondrial membrane, thereby clarifying, at the physical level, the spatial basis for its specific enrichment in mitochondrial-associated endoplasmic reticulum (MAMs) domains[38]. Further protein-protein interaction (PPI) network analysis revealed that ACSL4 exhibits close functional coupling with core anchoring proteins of MAMs (MFN2, CANX) and classical calcium channel complexes (VDAC1, HSPA9/GRP75). We further performed protein-protein docking simulations on the three-dimensional structures of IP_3_R and ACSL4. The results show that the two proteins may establish stable hydrogen bond and salt bridge structures at their interface via key amino acids (such as Cys595 and Glu666), with a weighted docking score of –1026 for the global minimum energy conformation. This provides physical confirmation that the core calcium channel and the lipid metabolic enzyme form an extremely compact spatial complex within the MAMs microdomain. This computational biology-based spatial coupling model was strongly corroborated by phenotypic evidence in our downstream functional experiments. Under the action of 3A72, not only were the expression levels of the core enzymes ACSL4 and LPCAT3, which mediate membrane lipid remodeling, significantly up-regulated, but the sharp increase in the concentration of the lipid peroxide MDA which is generated by their catalytic activity and the oxidative fluorescence of the C11-BODIPY probe also indicated that lipid peroxidation was occurring within the cells at this time.

At the ultrastructural level, 3A72 may precisely target and regulate the IP_3_R calcium channel through strong hydrophobic interactions with VAL794, THR795, and PRO796. Given the extremely high spatial topological proximity of IP_3_R and ACSL4 within MAMs, the sharp increase in local calcium flux induced by 3A72’s targeting of IP_3_R directly activates the neighboring ACSL4 metabolic hub as a key upstream signal. This cascade of calcium overload and lipid remodeling, driven by a spatial multi-anchor mechanism, leads to the vigorous accumulation of membrane lipid peroxides, fundamentally elucidating the molecular mechanism by which 3A72 specifically activates the MAMs structure to induce ferroptosis in TNBC cells[26].

This mechanistic model has also been comprehensively validated at the cellular phenotypic level. Experimental results show that 3A72 intervention not only leads to a significant upregulation of the ferroptosis-driving protein ACSL4 and depletion of the defense core GPX4, but also directly triggers the collapse of the antioxidant system and the massive accumulation of intracellular Fe^2^□.

In summary, our experimental results demonstrate that 3A72, through the release of NO and the activation of the chimeric IP_3_R/ACSL4 complex, induces sustained calcium overload, leading to lethal membrane phospholipid remodeling. Ultimately, this results in the specific induction of ferroptosis in TNBC cells via a cascade of lipid peroxidation triggered by calcium imbalance. However, to fully elucidate the pharmacological mechanism of action of 3A72, further refinement of our current work is required. In subsequent studies, we will endeavor to analyze in greater depth the precise targeting and microenvironmental effects of 3A72. Firstly, regarding the specific mechanism by which IP_3_R undergoes S-nitrosylation[39], current understanding remains largely based on theoretical predictions. Although molecular docking analyses have confirmed that the 3A72 molecule can bind to IP_3_R, there remains a considerable spatial distance (approximately 23.8Å) between the drug molecule and the predicted key cysteine residue. This implies that static docking models are insufficient to fully elucidate the dynamic modification process[40]. Consequently, the specific modification site and the actual binding mode still require definitive biochemical confirmation in future studies, utilizing the biotin-switch assay in conjunction with site-directed mutagenesis[41]. Secondly, given that 3A72 possesses both NO-donor properties and protein-binding activity, the relative contributions of 3A72’s release of free NO and its direct binding to IP_3_R/ACSL4 in inducing cellular oxidative stress have not yet been precisely quantified within the current experimental system. Finally, although this study has confirmed the cascade activation of the aforementioned key signaling axes and the expression dynamics of target proteins in an *in vivo* model, given the profound heterogeneity of the solid tumor microenvironment, The dynamic evolutionary process by which 3A72 in situ drives specific modifications of IP_3_R/ACSL4 requires further multidimensional and detailed characterization using intravital two-photon imaging[42]and specific probe tracing[43], in conjunction with spatial metabolomics[44].

In summary, this study elucidates the molecular mechanism by which the novel NO donor drug 3A72 induces ferroptosis in TNBC cells: not only does it cause cytosolic calcium overload and subsequent oxidative stress via NO release, but it also reshapes MAMs homeostasis and drives lipid peroxidation by precisely binding to MAMs-associated proteins IP_3_R and ACSL4. For the first time, a dual-pathway anti-tumor intervention strategy has been proposed for 3A72, combining the free release of NO with direct binding to targets. This study not only deepens our understanding of the pharmacological mechanisms underlying ferroptosis in TNBC but also provides a highly promising lead scaffold and translational basis for the future development of MAMs precision therapeutics.

## Author Contributions

YL (Yunke Long): Conceptualization, Investigation, Methodology, Data curation, Formal analysis, Writing original draft.

YJ (Yuxin Jiang): Data curation, Validation.

LL (La Li): Chemical Synthesis.

XW (Xiufan Wu): Data Analysis.

JW (Jiachen Weng): Data Analysis.

*YC (Ying Chen): Resources, Writing revision, Supervision.

*HL (Hongrui Liu): Conceptualization, Funding acquisition, Project administration, Supervision, Writing revision.

## Data Availability

The datasets generated and analyzed during the current study are kept confidential to protect intellectual property. However, raw data can be provided by the corresponding author to the journal’s editorial office upon request.

## Acknowledgement

This work is supported by the Scientific Research Innovation Program of the Shanghai Municipal Education Commission (No: 2023FGS2). We sincerely thank Servier Medical Art (http://smart.servier.com) for providing the elements of the graphical abstract. Furthermore, the results published here are in whole or part based upon data generated by the TCGA Research Network: https://www.cancer.gov/tcga. Molecular graphics and analyses performed with UCSF ChimeraX, developed by the Resource for Biocomputing, Visualization, and Informatics at the University of California, San Francisco, with support from National Institutes of Health R01-GM129325 and the Office of Cyber Infrastructure and Computational Biology, National Institute of Allergy and Infectious Diseases.

---

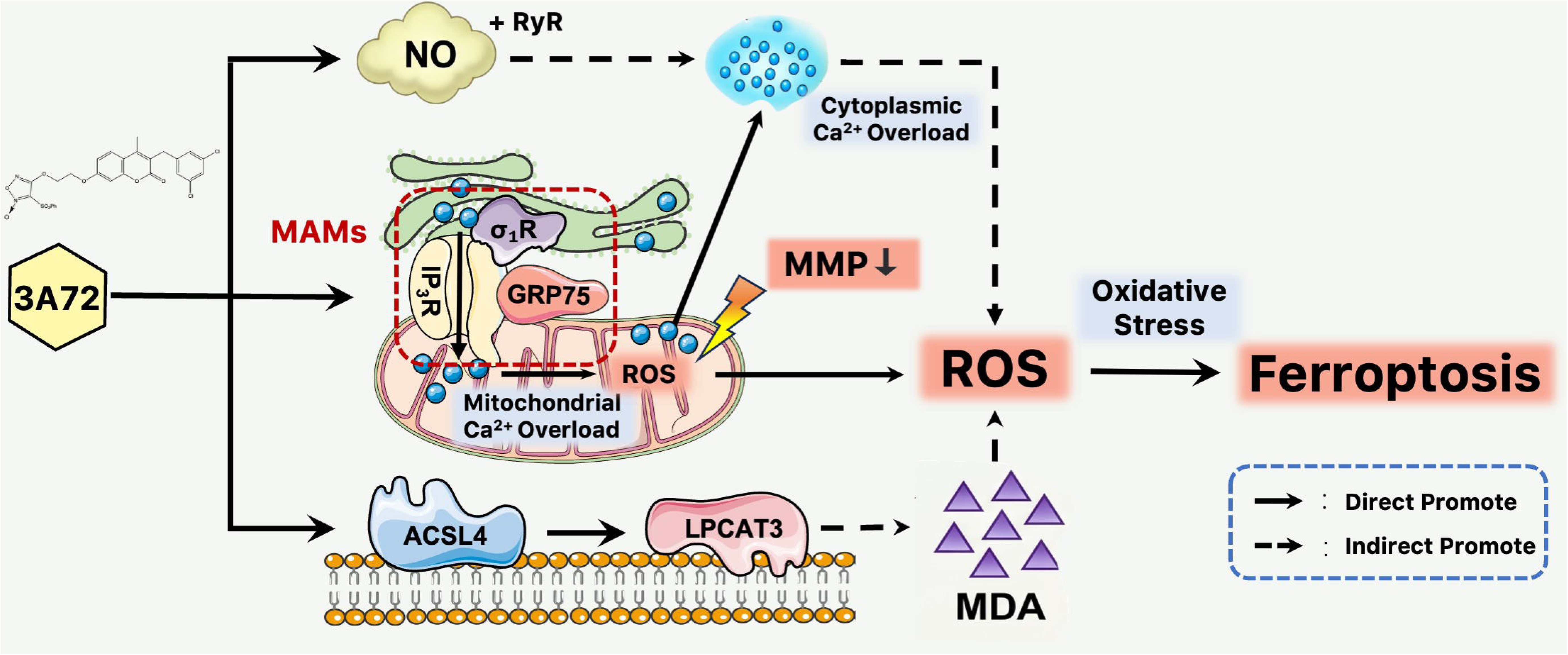

## Reference

[1] F. Derakhshan, J.S. Reis-Filho, Pathogenesis of Triple-Negative Breast Cancer, Annu. Rev. Pathol. Mech. Dis. 17 (2022) 181–204. 10.1146/annurev-pathol-042420-093238.

[2] L. Yin, J.-J. Duan, X.-W. Bian, S. Yu, Triple-negative breast cancer molecular subtyping and treatment progress, Breast Cancer Res 22 (2020) 61. 10.1186/s13058-020-01296-5.

[3] H. Li, P. Yang, J. Wang, J. Zhang, Q. Ma, Y. Jiang, Y. Wu, T. Han, D. Xiang, HLF regulates ferroptosis, development and chemoresistance of triple-negative breast cancer by activating tumor cell-macrophage crosstalk, J Hematol Oncol 15 (2022) 2. 10.1186/s13045-021-01223-x.

[4] A. Bardia, I.E. Krop, T. Kogawa, D. Juric, A.W. Tolcher, E.P. Hamilton, T. Mukohara, A. Lisberg, T. Shimizu, A.I. Spira, J. Tsurutani, S. Damodaran, K.P. Papadopoulos, J. Greenberg, F. Kobayashi, H. Zebger-Gong, R. Wong, Y. Kawasaki, T. Nakamura, F. Meric-Bernstam, Datopotamab Deruxtecan in Advanced or Metastatic HR+/HER2– and Triple-Negative Breast Cancer: Results From the Phase I TROPION-PanTumor01 Study, JCO 42 (2024) 2281–2294. 10.1200/JCO.23.01909.

[5] M. Damelin, A. Bankovich, J. Bernstein, J. Lucas, L. Chen, S. Williams, A. Park, J. Aguilar, E. Ernstoff, M. Charati, R. Dushin, M. Aujay, C. Lee, H. Ramoth, M. Milton, J. Hampl, S. Lazetic, V. Pulito, E. Rosfjord, Y. Sun, L. King, F. Barletta, A. Betts, M. Guffroy, H. Falahatpisheh, C.J. O’Donnell, R. Stull, M. Pysz, P. Escarpe, D. Liu, O. Foord, H.P. Gerber, P. Sapra, S.J. Dylla, A PTK7-targeted antibody-drug conjugate reduces tumor-initiating cells and induces sustained tumor regressions, Sci. Transl. Med. 9 (2017) eaag2611. 10.1126/scitranslmed.aag2611.

[6] Y. Li, H. Zhang, Y. Merkher, L. Chen, N. Liu, S. Leonov, Y. Chen, Recent advances in therapeutic strategies for triple-negative breast cancer, J Hematol Oncol 15 (2022) 121. 10.1186/s13045-022-01341-0.

[7] D.V. Ziegler, K. Parashar, L. Leal-Esteban, J. López-Alcalá, W. Castro, N. Zanou, L. Martinez-Carreres, K. Huber, X.P. Berney, M.M. Malagón, C. Roger, M.-A. Berger, Y. Gouriou, G. Paone, H. Gallart-Ayala, G. Sflomos, C. Ronchi, J. Ivanisevic, C. Brisken, J. Rieusset, M. Irving, L. Fajas, CDK4 inactivation inhibits apoptosis via mitochondria-ER contact remodeling in triple-negative breast cancer, Nat Commun 16 (2025) 541. 10.1038/s41467-024-55605-z.

[8] A.N. Von Krusenstiern, R.N. Robson, N. Qian, B. Qiu, F. Hu, E. Reznik, N. Smith, F. Zandkarimi, V.M. Estes, M. Dupont, T. Hirschhorn, M.S. Shchepinov, W. Min, K.A. Woerpel, B.R. Stockwell, Identification of essential sites of lipid peroxidation in ferroptosis, Nat Chem Biol 19 (2023) 719–730. 10.1038/s41589-022-01249-3.

[9] F. Barberini, R. Pietroni, S. Ielpo, V. Lucarini, D. Nardozi, O. Melaiu, M. Benvenuto, C. Focaccetti, C. Palumbo, F. Rossin, D. Fruci, D. Olive, L. Masuelli, R. Bei, L. Cifaldi, Combined IFN-γ and TNF-α treatment enhances the susceptibility of breast cancer cells and spheroids to Natural Killer cell-mediated killing, Cell Death Dis 16 (2025) 729. 10.1038/s41419-025-08021-0.

[10] W. Zhao, R. Sheng, The correlation between mitochondria-associated endoplasmic reticulum membranes (MAMs) and Ca2+ transport in the pathogenesis of diseases, Acta Pharmacol Sin 46 (2025) 271–291. 10.1038/s41401-024-01359-9.

[11] X. Zhao, L. Yan, L. Tian, X. Zhang, R. Liu, Z. Li, Metformin improves intestinal ischemia-reperfusion injury by reducing the formation of mitochondrial associated endoplasmic reticulum membranes (MAMs) and inhibiting ferroptosis in intestinal cells, Front. Pharmacol. 16 (2025) 1581085. 10.3389/fphar.2025.1581085.

[12] H. Liu, S. Zheng, G. Hou, J. Dai, Y. Zhao, F. Yang, Z. Xiang, W. Zhang, X. Wang, Y. Gong, L. Li, N. Zhang, Y. Hu, AKAP1/PKA-mediated GRP75 phosphorylation at mitochondria-associated endoplasmic reticulum membranes protects cancer cells against ferroptosis, Cell Death Differ 32 (2025) 488–505. 10.1038/s41418-024-01414-2.

[13] R. Fu, L. Fang, J. Liu, Y. Liu, Y. Wang, D. Kong, J. Guo, Mitochondrial-associated endoplasmic reticulum membranes (MAMs) drive ferroptosis via redox signaling: A key potential mechanism in epilepsy, Neurobiology of Disease 218 (2026) 107248. 10.1016/j.nbd.2025.107248.

[14] W. Fernando, J. Clucas, A. Rizzo, R. Singer, E. Goode, C. Tiu, S. Layzell, J. Konecnik, R. Wilson, S. Wicky John, S. Jouny, N. Guppy, V. Boulat, J. Mannion, M. Goicoechea, S. Tan, S. Lawson, C. Starling, G. Elshtein, N. Ravindran, A.B. Montgomery, R. Andres-Ejarque, M. Allen, S. Lumbard, F. Wallberg, K. Betteridge, R. Scrimgeour, D. Robertson, G. Ward, M. Sims, T. Smyth, A.L. Samson, J.M. Murphy, D. Kolarevic Ivankovic, E.N. Arwert, D.P. Calado, A. Grigoriadis, M.J. Smalley, A. Melcher, S. Haider, T. Lawrence, I. Roxanis, A.N.J. Tutt, T. Tenev, P. Meier, Necroptosis in both tumour and stromal compartments determines responsiveness to immunogenic cell death-based immunotherapy, Nat Commun 17 (2026) 3597. 10.1038/s41467-026-70133-8.

[15] J. Lu, Y. Feng, Y. Wang, Y. Yu, W. Zhao, X. Cao, Z. Li, Y. Lu, Nitric oxide induces apoptosis of human primary melanocytes by regulating calcium homeostasis via VDAC1, Mol Cell Biochem 480 (2025) 6111–6125. 10.1007/s11010-025-05361-5.

[16] S. Xin, C. Mueller, S. Pfeiffer, V.A.N. Kraft, J. Merl-Pham, X. Bao, R. Feederle, X. Jin, S.M. Hauck, P. Schmitt-Kopplin, J.A. Schick, MS4A15 drives ferroptosis resistance through calcium-restricted lipid remodeling, Cell Death Differ 29 (2022) 670–686. 10.1038/s41418-021-00883-z.

[17] D.S. Chandrashekar, S.K. Karthikeyan, P.K. Korla, H. Patel, A.R. Shovon, M. Athar, G.J. Netto, Z.S. Qin, S. Kumar, U. Manne, C.J. Creighton, S. Varambally, UALCAN: An update to the integrated cancer data analysis platform, Neoplasia 25 (2022) 18–27. 10.1016/j.neo.2022.01.001.

[18] D.S. Chandrashekar, B. Bashel, S.A.H. Balasubramanya, C.J. Creighton, I. Ponce-Rodriguez, B.V.S.K. Chakravarthi, S. Varambally, UALCAN: A Portal for Facilitating Tumor Subgroup Gene Expression and Survival Analyses, Neoplasia 19 (2017) 649–658. 10.1016/j.neo.2017.05.002.

[19] S.V. Vasaikar, P. Straub, J. Wang, B. Zhang, LinkedOmics: analyzing multi-omics data within and across 32 cancer types, Nucleic Acids Research 46 (2018) D956– D963. 10.1093/nar/gkx1090.

[20] D. Szklarczyk, R. Kirsch, M. Koutrouli, K. Nastou, F. Mehryary, R. Hachilif, A.L. Gable, T. Fang, N.T. Doncheva, S. Pyysalo, P. Bork, L.J. Jensen, C. von Mering, The STRING database in 2023: protein–protein association networks and functional enrichment analyses for any sequenced genome of interest, Nucleic Acids Research 51 (2023) D638–D646. 10.1093/nar/gkac1000.

[21] X. Yang, Y. Liu, J. Gan, Z.-X. Xiao, Y. Cao, FitDock: protein–ligand docking by template fitting, Briefings in Bioinformatics 23 (2022) bbac087. 10.1093/bib/bbac087.

[22] Y. Liu, X. Yang, J. Gan, S. Chen, Z.-X. Xiao, Y. Cao, CB-Dock2: improved protein– ligand blind docking by integrating cavity detection, docking and homologous template fitting, Nucleic Acids Research 50 (2022) W159–W164. 10.1093/nar/gkac394.

[23] G. Jones, A. Jindal, U. Ghani, S. Kotelnikov, M. Egbert, N. Hashemi, S. Vajda, D. Padhorny, D. Kozakov, Elucidation of protein function using computational docking and hotspot analysis by ClusPro and FTMap, Acta Crystallogr D Struct Biol 78 (2022) 690–697. 10.1107/S2059798322002741.

[24] X. Zhou, C. Gao, C. Liu, R. Zhang, Z. Hu, U. Ali, L. Zhang, X. Chen, B. Liu, L. Li, C. Wang, Regulating cell stress responses by nitric oxide to enhance calcium overload-mediated tumor therapy, Biomaterials 326 (2026) 123676. 10.1016/j.biomaterials.2025.123676.

[25] Y. Gong, X. Lu, X. Wang, Y. Wang, Z. Shen, Y. Gao, L. Zhuang, L. Yu, J. Chen, Q. Li, F. Qiu, J. Lin, Y. Tao, C. Jiang, G. Fu, P. Zhang, D. Lai, Mitochondrial Tumor Suppressor 1A Attenuates Myocardial Infarction Injury by Maintaining the Coupling Between Mitochondria and Endoplasmic Reticulum, Circulation 152 (2025) 183–201. 10.1161/CIRCULATIONAHA.124.069737.

[26] D. Liang, Y. Feng, F. Zandkarimi, H. Wang, Z. Zhang, J. Kim, Y. Cai, W. Gu, B.R. Stockwell, X. Jiang, Ferroptosis surveillance independent of GPX4 and differentially regulated by sex hormones, Cell 186 (2023) 2748–2764.e22. 10.1016/j.cell.2023.05.003.

[27] V. Manganelli, P. Matarrese, M. Antonioli, L. Gambardella, T. Vescovo, C. Gretzmeier, A. Longo, A. Capozzi, S. Recalchi, G. Riitano, R. Misasi, J. Dengjel, W. Malorni, G.M. Fimia, M. Sorice, T. Garofalo, Raft-like lipid microdomains drive autophagy initiation via AMBRA1-ERLIN1 molecular association within MAMs, Autophagy 17 (2021) 2528–2548. 10.1080/15548627.2020.1834207.

[28] Z. Wang, Z. Li, L. Wang, H. Zhang, X. Yang, L. Bao, G. Dou, L. Ren, Y. Fu, L. Li, S. Gong, Y. Zhou, F. Ding, L. Yu, H. Luo, Y. Liu, F. Zhang, H. Yu, S. Liu, X. Liu, F. Wei, S. Liu, Cellular Adaptation to Mechanical Stress Emerges via Cell Shrinkage Triggered by Nonlinear Calcium Elevation, Advanced Science 12 (2025) e03659. 10.1002/advs.202503659.

[29] J. Li, F. Qi, H. Su, C. Zhang, Q. Zhang, Y. Chen, P. Chen, L. Su, Y. Chen, Y. Yang, Z. Chen, S. Zhang, GRP75-faciliated Mitochondria-associated ER Membrane (MAM) Integrity controls Cisplatin-resistance in Ovarian Cancer Patients, Int. J. Biol. Sci. 18 (2022) 2914–2931. 10.7150/ijbs.71571.

[30] Y. Qiu, X. Wang, Y. Sun, T. Jin, R. Tang, X. Zhou, M. Xu, Y. Gan, R. Wang, H. Luo, M. Liu, X. Tang, ACSL4-Mediated Membrane Phospholipid Remodeling Induces Integrin β1 Activation to Facilitate Triple-Negative Breast Cancer Metastasis, Cancer Research 84 (2024) 1856–1871. 10.1158/0008-5472.CAN-23-2491.

[31] K.-J. Chen, Y. Zhang, X.-Y. Zhu, S. Yu, Y. Xie, C.-J. Jin, Y.-M. Shen, S.-Y. Zhou, X.-C. Dai, S.-A. Su, L. Xie, Z.-X. Huang, H. Gong, M.-X. Xiang, H. Ma, GSTM1 suppresses cardiac fibrosis post-myocardial infarction through inhibiting lipid peroxidation and ferroptosis, Military Med Res 12 (2025) 26. 10.1186/s40779-025-00610-6.

[32] A.F. Krüger, W. Schmitz, S. Lamer, A. Triebig, T. Maier, C.T. Fuss, J.P.F. Angeli, A. Schlosser, C. Stigloher, M. Fassnacht, I. Weigand, M. Kroiss, Adrenocortical Mitochondria-Associated Membranes: Isolation, Characterization, and Lipidoproteomic Response to Mitotane, Journal of the Endocrine Society 10 (2025) bvaf198. 10.1210/jendso/bvaf198.

[33] D. Xiao, W. Chang, X. Ao, L. Ye, W. Wu, L. Song, X. Yuan, L. Feng, P. Wang, Y. Wang, Y. Jia, X. Tang, J. Wang, Parkin inhibits iron overload-induced cardiomyocyte ferroptosis by ubiquitinating ACSL4 and modulating PUFA-phospholipids metabolism, Acta Pharmaceutica Sinica B 15 (2025) 1589–1607. 10.1016/j.apsb.2024.12.027.

[34] M.-D. Li, L. Fu, B.-B. Lv, Y. Xiang, H.-X. Xiang, D.-X. Xu, H. Zhao, Arsenic induces ferroptosis and acute lung injury through mtROS-mediated mitochondria-associated endoplasmic reticulum membrane dysfunction, Ecotoxicology and Environmental Safety 238 (2022) 113595. 10.1016/j.ecoenv.2022.113595.

[35] K. Ooi, L. Hu, Y. Feng, C. Han, X. Ren, X. Qian, H. Huang, S. Chen, Q. Shi, H. Lin, J. Wang, D. Zhu, R. Wang, C. Xia, Sigma-1 Receptor Activation Suppresses Microglia M1 Polarization via Regulating Endoplasmic Reticulum–Mitochondria Contact and Mitochondrial Functions in Stress-Induced Hypertension Rats, Mol Neurobiol 58 (2021) 6625–6646. 10.1007/s12035-021-02488-6.

[36] Q. He, M. Qu, T. Shen, J. Su, Y. Xu, C. Xu, M.Q. Barkat, J. Cai, H. Zhu, L.-H. Zeng, X. Wu, Control of mitochondria-associated endoplasmic reticulum membranes by protein S-palmitoylation: Novel therapeutic targets for neurodegenerative diseases, Ageing Research Reviews 87 (2023) 101920.10.1016/j.arr.2023.101920.

[37] A.M. Karim, J. Eun Kwon, T. Ali, J. Jang, I. Ullah, Y.-G. Lee, D.W. Park, J. Park, J.W. Jeang, S.C. Kang, Triple-negative breast cancer: epidemiology, molecular mechanisms, and modern vaccine-based treatment strategies, Biochemical Pharmacology 212 (2023) 115545. 10.1016/j.bcp.2023.115545.

[38] J. Lu, Y. Tao, J. Li, M. Wubuliaila, J. Wang, N. Li, TP53 links mitochondria– endoplasmic reticulum crosstalk and ferroptosis in septic cardiomyopathy: Protective modulation by nicorandil, Cellular Signalling 144 (2026) 112537. 10.1016/j.cellsig.2026.112537.

[39] S.Y. Park, K. Zhu, Advances in SnO_2_ for Efficient and Stable n–i–p Perovskite Solar Cells, Advanced Materials 34 (2022) 2110438. 10.1002/adma.202110438.

[40] P.-T. Doulias, J.L. Greene, T.M. Greco, M. Tenopoulou, S.H. Seeholzer, R.L. Dunbrack, H. Ischiropoulos, Structural profiling of endogenous S-nitrosocysteine residues reveals unique features that accommodate diverse mechanisms for protein S-nitrosylation, Proc. Natl. Acad. Sci. U.S.A. 107 (2010) 16958–16963. 10.1073/pnas.1008036107.

[41] H.S. Chung, C.I. Murray, V. Venkatraman, E.L. Crowgey, P.P. Rainer, R.N. Cole, R.D. Bomgarden, J.C. Rogers, W. Balkan, J.M. Hare, D.A. Kass, J.E. Van Eyk, Dual Labeling Biotin Switch Assay to Reduce Bias Derived From Different Cysteine Subpopulations: A Method to Maximize S-Nitrosylation Detection, Circulation Research 117 (2015) 846–857. 10.1161/CIRCRESAHA.115.307336.

[42] Z. Zhao, Y. Zhou, B. Liu, J. He, J. Zhao, Y. Cai, J. Fan, X. Li, Z. Wang, Z. Lu, J. Wu, H. Qi, Q. Dai, Two-photon synthetic aperture microscopy for minimally invasive fast 3D imaging of native subcellular behaviors in deep tissue, Cell 186 (2023) 2475–2491.e22. 10.1016/j.cell.2023.04.016.

[43] K.R. Mesa, K.A. O’Connor, C. Ng, S.P. Salvatore, A. Dolynuk, M.R. Lomeli, D.R. Littman, Niche-specific dermal macrophage loss promotes skin capillary ageing, Nature 648 (2025) 173–181. 10.1038/s41586-025-09639-y.

[44] J. Zhou, H. Zhang, W.-W. Pan, C. Peng, H. Tang, G. Yuan, F. Peng, Isocucurbitacin B targets STAT3 to induce ferroptosis and promote anti-PD1 immunotherapy responses in breast cancer, International Journal of Surgery 112 (2026) 11404– 11424. 10.1097/JS9.0000000000004907.

